# A curcumin direct protein (DiPro) biosensor for cell-free prototyping

**DOI:** 10.1101/2021.09.22.461347

**Authors:** Agata Lesniewska, Guy Griffin, Paul S Freemont, Karen M Polizzi, Simon J Moore

## Abstract

In synthetic biology, biosensors are routinely coupled to a gene expression cascade for detecting small molecules and physical signals. We posit that an alternative direct protein (DiPro) biosensor mechanism, could provide a new opportunity for rapid detection of specific chemicals. Herein, we reveal a fluorescent curcumin DiPro biosensor, based on the *Escherichia* coli double bond reductase (*Ec*CurA) as a detection system. We characterise the *Ec*CurA DiPro biosensor and propose enhanced curcumin fluorescence is generated through π-π stacking between protein and ligand. Using a cell-free synthetic biology approach, we use the *Ec*CurA DiPro biosensor to fine tune 10 reaction parameters (cofactor, substrate, and enzyme levels) for cell-free biosynthesis, assisted through acoustic liquid handling robotics. Overall, we increase *Ec*CurA-curcumin fluorescence by 80-fold. We speculate that a generic DiPro biosensor fluorescence mechanism can be further exploited for a wider range of chemicals that share intrinsic fluorescence and have a suitable binding protein.

## Introduction

There is a rising interest to develop biosensors that can detect natural products and industrial chemicals, in order to engineer heterologous production in microbial/plant cells and cell-free systems^1,2^. This is important, as most chemicals require resource-intense analytical processes (e.g., high-performance liquid chromatography, mass spectrometry) for quantitation^3^. This type of analysis presents a bottleneck for the implementation of the design-build-test-learn (DBTL) cycle within synthetic biology^1^. Biosensors including riboswitches and transcription factors (TFs) are routinely coupled to a fluorescence protein^4–6^, RNA aptamer^7^ or enzymatic reaction^8,9^, to control gene expression. Transcription factors (TFs) are the largest family of biosensors and use either a one- or two-component cascade^10–12^, to sense a range of metabolites and physical signals. About 4,000 TFs are listed on the PRODORIC2® database currently^12^. In contrast, riboswitches are *cis* RNA elements that form secondary structures to interact with target ligands and regulate either transcription or translation. Riboswitches can detect some amino acids, metals, nucleotides and cofactors^13^. However, both riboswitches and TFs require a gene expression cascade to regulate an output device (e.g., fluorescence protein). Therefore, their sensory function can be described as indirect, complicating the measurement workflow.

We posited that an alternative yet generalisable direct biosensor approach exists where the signal is obtained from direct interactions of the analyte with the biosensor. Such a mechanism bypasses the typical gene expression/maturation requirements of oxygen-dependent fluorescent proteins in genetic circuit design^14^. In addition, it is desirable to target industrially relevant chemicals (e.g., natural products, fine chemicals) outside of the typical substrate scope of TFs (primary metabolites/physical stimuli) and riboswitches. Therefore, we reviewed the literature for potential untapped sources of direct biosensors, such as fluorescent protein-ligand complexes; whereby fluorescence provides a molecular handle to gauge ligand concentration. Examples, though few, are led by nucleotide-binding proteins, such as flavin mononucleotide (FMN)-binding Lov proteins^15^, blue-fluorescent NADPH-oxidoreductase^16^ and the NAD^+^/NADH binding Rex protein^17^, albeit fused to a circularly permuted yellow fluorescence protein (YFP) for detection^18^. Interestingly, these interactions exploit the intrinsic fluorescence of the nucleotides FMN and NAD(P)H. An exception to this nucleotide biosensor collection, is the biliverdin binding small ultra-red fluorescent protein (smURFP)^19^. At this point, we speculated whether other small molecules that possess similar chemical properties as biliverdin (e.g., conjugation, aromatic), and intrinsic fluorescence, could provide new biosensor leads. Hereafter, we collectively refer to such exemplars as direct protein (DiPro) biosensors.

## Results and Discussion

Herein, we report a DiPro biosensor, based on the intrinsic fluorescence properties of the natural product curcumin. Curcumin is a type III polyketide and a yellow pigment produced by the turmeric plant (*Curcuma longa*). From a biotechnology perspective, there is a generic interest in using microbes to make curcumin, a medicinal natural product, in order to replace land-intense farming of turmeric^20^. Curcumin biosynthesis stems from L-tyrosine/L-phenylalanine metabolism. The molecule displays weak intrinsic fluorescence in aqueous solution with a quantum yield of 0.01^21^. However, curcumin fluorescence is strongly enhanced upon non-specific binding to extracellular curli (amyloid-like fibres) on the surface of *E. coli*^22^ – a physical property that we sought to exploit. Next, we investigated the curcumin binding properties of the NADPH-dependent *E. coli* curcumin reductase (*Ec*CurA - NCBI RefSeq: WP_000531452), as a potential soluble intracellular protein that may mirror the curcumin-curli fluorescence interaction. To begin, we constitutively expressed and purified N-terminal His_6_-tagged *Ec*CurA to homogeneity. A single litre of *E. coli* BL21 (DE3) Star culture in 2YT provided approximately 20 mg of pure apoprotein with an observed molecular weight of ∼40 kDa (His_6_-tag-*Ec*CurA is 39.6 kDa) by denaturing gel electrophoresis (**Figure 1A**). Favourably, *Ec*CurA accumulated about ∼30% of total protein content in *E. coli* and was located entirely in the soluble fraction. Post-purification, *Ec*CurA was highly stable up to 30 mg/mL and could be stored in the freezer without cryoprotectants (e.g., glycerol, DMSO). In addition, we also incubated an excess of curcumin with the cell-extract and purified (see methods). This resulted in a visual co-purification of *Ec*CurA in complex with curcumin. Curcumin displayed an absorbance maximum at 425 nm (ε = 23,800 M^-1^ cm^-1^) in aqueous solution^23^. In contrast, UV-Visible absorbance spectra of the co-purified *Ec*CurA-curcumin complex was blue-shifted to 384 nm for the curcumin absorbance maxima (**Figure 1B**). Next, we incubated 25 μM of the apoprotein with 25 μM of curcumin. Binding of curcumin to *Ec*CurA generated strong yellow fluorescence, with broad absorption and emission spectra with maxima at 427 nm and 522 nm, respectively (**Figure 1C**). Fluorescence was relatively stable up to an hour, although curcumin is known to spontaneously degrade in alkaline conditions^21^. Since *Ec*CurA has catalytic activity as a double bond reductase, the addition of an excess of NADPH (1 mM) led to substrate turnover, resulting in a loss of conjugation and visual colour/fluorescence. While previous studies on the *Ec*CurA and related *Vibrio vulnificus* CurA (*Vv*CurA) have established the kinetics of CurA^24,25^, both enzymes were studied at sub-stoichiometric levels. In contrast, by incubating 25 μM of *Ec*CurA with a titration of curcumin (0-100 μM), the fluorescence signal fitted an exponential curve with a *K*_D_ of 8.39 μM (**Figure S1-S3**). In addition, the *Ec*CurA DiPro biosensor showed a reasonable limit of detection of 0.29 μM for curcumin, calculated as previously described^26^. Regarding the fluorescence mechanism of the *Ec*CurA DiPro biosensor, the predicted active site is highly aromatic binding pocket (Y53, Y64 and Y253) near the NADPH binding cavity. During our study, the highly related *Vv*CurA structure (PDB: 5ZXN) was released with NADP^+^ bound^24^. We compared *Vv*CurA to our own related reductase structure (PDB: 6EOW) with NADP+ and half-curcumin (*p*-hydroxybenzalacetone) bound^27^. Based on this, we speculate that curcumin fluorescence (in the absence of NADPH), is generated through stabilization of π-π* transitions upon excitation (450 nm), leading to an excited state intramolecular charge transfer species^21^. Enhanced fluorescence is likely due to interactions between one-half of the curcumin molecule (the benzalacetone moiety), and the aromatic binding pocket of *Ec*CurA^21^.

**Figure 1.**
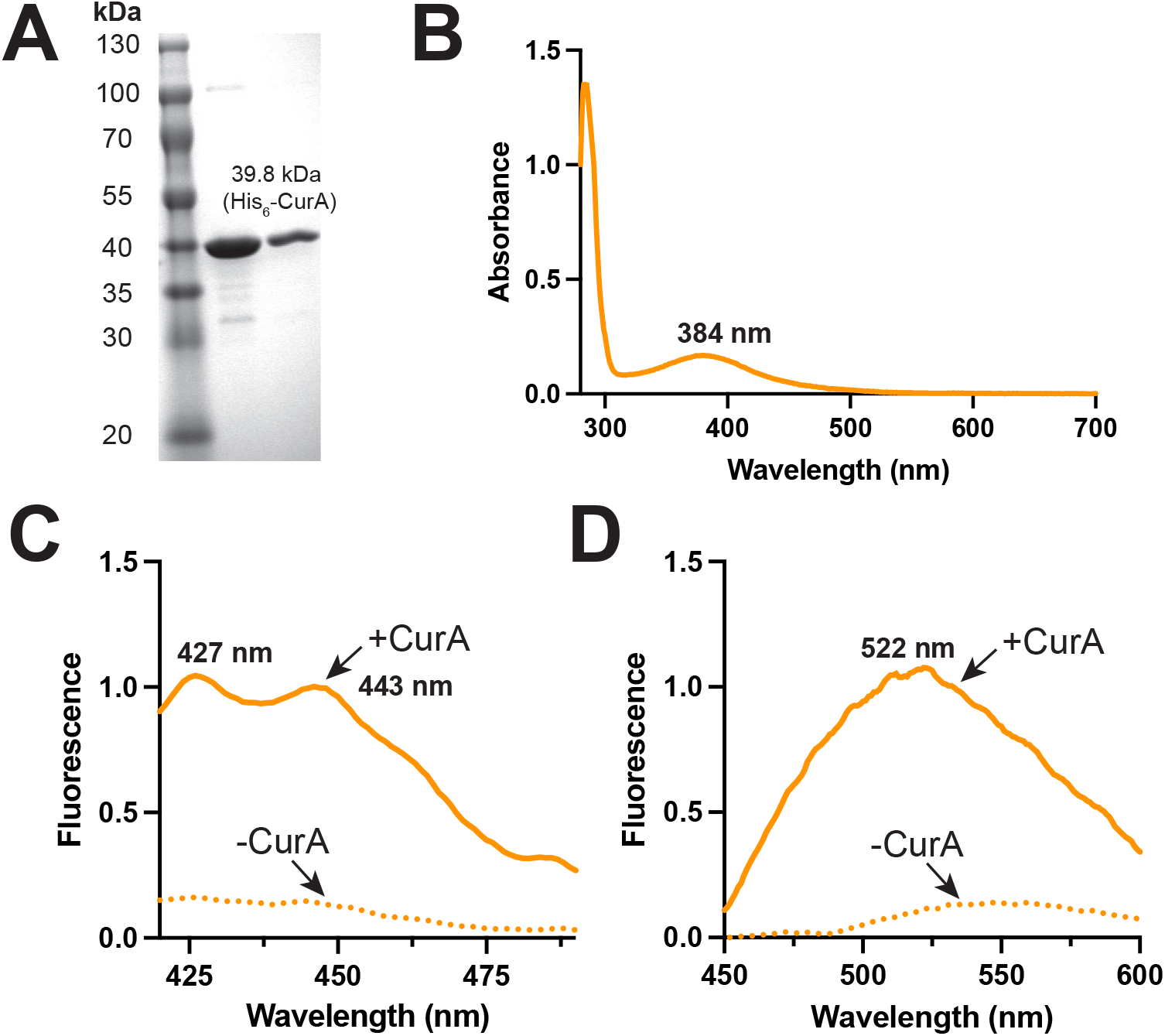
Characterisation of the *Ec*CurA DiPro biosensor. (A) SDS-PAGE of purified His_6_-tagged *Ec*CurA. (B) UV-Visible absorbance of co-purified His_6_-tagged *Ec*CurA bound with curcumin. (C) Relative excitation and (D) emission fluorescence spectra of *Ec*CurA with and without curcumin bound (1:1 ratio – 25 μM) in Buffer A at 25°C.

Next, we sought to demonstrate the potential of the *Ec*CurA DiPro biosensor using a cell-free synthetic biology strategy. Here we added an excess of the *Ec*CurA DiPro biosensor in a purified multienzyme system in the absence of NADPH. Curcumin biosynthesis, like other type III polyketides, requires the precursors *p*-coumaroyl-CoA (and analogs) and malonyl-CoA. We therefore created a synthetic enzyme pathway to supply these precursors and study the biosynthesis of curcumin. For this we used the tyrosine ammonia lyase (TAL), *p*-coumaroyl-CoA ligase and malonyl-CoA synthetase (MatB), described in our previous study^27^, along with a curcumin synthase (CUS) from *Oryza sativa*^29^. Using 1 mM L-tyrosine as a substrate, the cofactors (ATP, malonate, CoA, Mg^2+^) and purified enzymes were included (see methods), to attempt to synthesize the analog of curcumin, bisdemethoxycurcumin (BDMC). Firstly, initial reactions accumulated a yellow visual appearance (**Figure 2A**). Critically, the reaction product was fluorescent only in the presence of the *Ec*CurA (25 μM) DiPro biosensor (**Figure 2A**) and we confirmed the biosynthesis of the product from the reaction using HPLC. Next, we prepared a time-course reaction to monitor BDMC biosynthesis using the *Ec*CurA DiPro biosensor as a detector. Here, increasing fluorescence was observed, relative to background. Importantly, if the *Ec*CurA DiPro biosensor or any of the biosynthetic enzymes or substrates were omitted, fluorescence dropped to background levels (**Figure S4**). Furthermore, if an aliquot of NADPH (1 μM) was spiked into the reaction, a drop in fluorescence was observed, showing that *Ec*CurA-dependent reductase activity quenches fluorescence. These results confirmed the sensitivity and specificity of the assay for BDMC biosynthesis.

**Figure 2.**
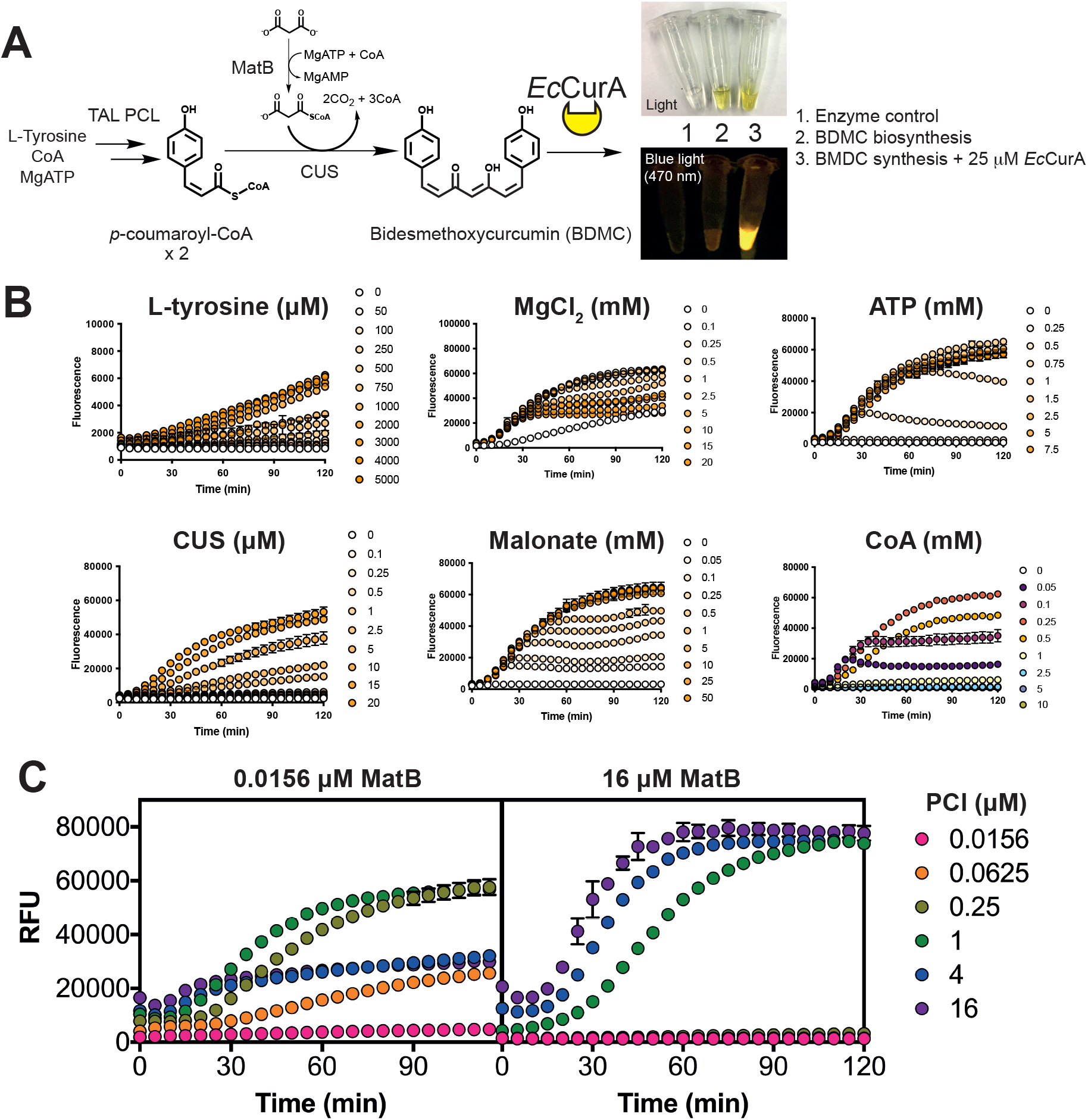
Optimisation curcumin biosynthesis with the DiPro biosensor, assisted with acoustic liquid handling robotics. (A) A synthetic pathway for detection of BDMC synthesis from malonyl-CoA and *p*-coumaroyl-CoA. *Ec*CurA binds to BDMC/curcumin generating a unique fluorescent output for relative quantitation of pathway activity. Inset visual image shows visible fluorescence of cell-free BDMC reactions, with negative controls. (B) Fine-tuning of cofactors and substrates, and (C) enzymes to maximise BDMC fluorescence signal detection with *Ec*CurA.

An important advantage for a cell-free multienzyme system, is the ability to rapidly optimise multiple parameters simultaneously. Therefore, to rapidly optimise curcumin biosynthesis, we used the *Ec*CurA DiPro biosensor to monitor multiple reactions in parallel using real-time fluorescence measurements. This was assisted with a pick-list generating script and an acoustic liquid handling robot, as described previously^30^. Firstly, L-tyrosine and CUS levels were set at 5 mM and 25 μM, respectively, so that these parameters were not rate limiting (**Figure 2B**). Next, BDMC biosynthesis was optimised, through *Ec*CurA DiPro biosensor fluorescence, with respect to ATP, Mg^2+^, CoA and malonate levels. Due to the non-linear relationship of fluorescence to curcumin/BDMC concentration, we report relative fluorescence units. We found that the ATP level was critical to *Ec*CurA DiPro biosensor fluorescence with at least 1 mM required to reach saturation (∼65,000 RFU), while for optimal malonyl-CoA supply, a concentration of greater than 2.5 mM malonate was required for maximal activity (>60,000 RFU) and fluorescence saturation. The Mg^2+^ level between 0.1 and 1 mM supported similar fluorescence levels at 0.1-1 mM, but with higher concentrations, a biphasic curve was observed (**Figure 2B**). A similar observation also occurred with the levels of CoA, where concentrations above or below the optimum CoA (0.25 mM) concentration. One potential interpretation of this response is a temporary depletion of free CoA availability, thus leading to an imbalance between *p*-coumaroyl-CoA and malonyl-CoA levels for the CUS enzyme. In contrast, if higher levels of CoA (>1 mM) were added, reduced *Ec*CurA DiPro biosensor fluorescence was observed (**Figure 2B**). We next used this optimised set of conditions to determine the optimal stoichiometry of enzymes in the biosynthetic pathway. To do this, each enzyme was varied in a 4-fold dilution series from 0.0156 to 16 μM. Firstly, some background fluorescence attributed to high levels of *p*-coumaroyl-CoA occurred if the level of the PCL enzyme was increased above 1 μM (**Figure 2C**). However, if the levels of the TAL and PCL enzymes were optimised to a peak concentration of 16 μM and 1 μM, respectively, maximal *Ec*CurA DiPro biosensor fluorescence (87,400 RFU) was reached, an 80-fold increase from starting conditions. Interestingly, by varying PCL and MatB together, a careful balance of enzymes was required to prevent pathway inhibition, which indicates rapid depletion of free CoA (**Figure 2C**).

In summary, our proof-of-concept *Ec*CurA DiPro biosensor permits rapid prototyping of a cell-free biosynthetic pathway for a curcumin analogue in microscale conditions. While our experiments only provide a relative measure of pathway activity, it demonstrates a finely tuned interplay between enzyme levels and the CoA cofactor is required for optimal pathway performance. This observation demonstrates a clear advantage of studying enzyme ensembles from a cell-free angle. Further work is required to engineer *Ec*CurA DiPro biosensor for cell-based application by eliminating NADPH binding or catalytic activity. Alternatively, the binding pocket could be engineered to target new high-value chemicals that interact and fluorescence with the aromatic core. In conclusion, our report highlights a potential new DiPro biosensor mechanism using a fluorescent protein-substrate binary complex. This approach is advantageous as input to output signal propagation is rapid; it does not require post gene expression or protein synthesis/maturation. We postulate that a wider variety of DiPro biosensors is within reach, particularly for pigmented natural products with intrinsic fluorescence. Current estimates suggest that approximately 300 natural products (not including synthetic chemicals and analogs) have some form of intrinsic fluorescence and therefore provide a putative target for future DiPro biosensor development^32^.

## Materials and Methods

### Molecular biology

Routine molecular biology was performed as described previously^33^. The TAL, PCL and MatB expression constructs were described in our previous work^27^. CUS^29^ was synthesised by ThermoFisher Scientific and codon optimised for *E. coli* K12 expression with compatibility for EcoFlex cloning^33^. *Ec*CurA was PCR amplified from *E. coli* MG1655 genomic DNA using Q5 polymerase and sub-cloned into pBP-ORF^33^. *Ec*CurA was assembled with a strong constitutive promoter (SJM928) and N-terminal His_6_-tag for *E. coli* expression. All oligonucleotides, plasmids and synthetic DNA sequences are listed in the supporting information. Sequencing was performed by Eurofins, UK.

### Protein expression and purification

His_6_-tagged recombinant TAL, PCL, CUS, MatB and *Ec*CurA were over-produced in *E. coli* BL21-Gold (DE3) grown at 37°C, 200 rpm in 2YT medium with 100 μg/mL ampicillin until an OD_600_ of 0.6 was reached. Cells were induced with 0.4 mM IPTG and grown overnight at 21°C at 200 rpm. Cell were collected by centrifugation at 6,000 × *g*, 4°C for 20 min, then re-suspended in binding buffer (20 mM Tris-HCl pH 8, 500 mM NaCl, 5 mM imidazole) and lysed by sonication. Cell-lysates were clarified with centrifugation at 45,000 × *g*, 4°C for 20 min and purified by gravity flow using Ni-nitrilotriacetic acid (NTA) agarose (Cytiva). His_6_-tagged proteins were washed with increasing concentrations of imidazole (5, 30 and 70 mM) in 20 mM Tris-HCl pH 8, 500 mM NaCl, before elution at 400 mM imidazole. Purified proteins were dialysed (MWCO 10,000) for ∼16 hours in 2 litres of 20 mM HEPES pH 7.5, 100 mM NaCl (Buffer A) at 4°C. All enzymes were soluble and retained activity upon storage at −80°C with 15% (v/v) glycerol. Aliquots of 20 mg/mL *Ec*CurA were stored in Buffer A without glycerol at − 80°C.

### Absorbance spectroscopy and fluorescence titrations

Absorbance and fluorescence spectra of purified *Ec*CurA and controls were measured in Buffer A, using an Agilent Cary60 UV-Vis or Cary Eclipse fluorescence spectrometer, respectively.

### Liquid handling robotics and platereader fluorescence measurements

Reactions were studied in 384 or 1536 well microtiter plates (Greiner), at a volume of 10 μL or 2 μL, respectively. Reactions were prepared using an Echo® 525 acoustic liquid handling robot (LabCyte). A general Python script for robot transfer instructions is available at https://github.com/jmacdona/ODE_MCMC_tools. Liquid droplets were transferred as multiples of 25 nL to a final volume of 2-10 μL as technical triplicate replicates. Plates were sealed with Breathe-Easy^®^ sealing membrane (Sigma) and briefly centrifuged at 1,000 × *g* for 10 seconds. A CLARIOStar plate reader (BMG Labtech, Germany) was used for enzyme incubations and kinetic fluorescence measurements. Standard measurements were as follows: 30°C incubation temperature, 40 flashes per well, 10 seconds of 300 rpm orbital shaking prior to measurement, 0.1 s settling time, 1 min recordings for 180 cycles, 425-10 nm excitation and 520-20 nm emission.

## Acknowledgements

SJM would like to acknowledge the following research support: EPSRC [EP/K038648/1] for SJM as a postdoc with PSF/KP; High-Value Biorenewables (BBSRC) network business-interaction voucher funding for SJM as principal investigator. SJM would also like to thank Professor Neil Kad (University of Kent) for discussions on the binding and fluorescence kinetics of *Ec*CurA.

## Supplementary Data

**Figure S1.**
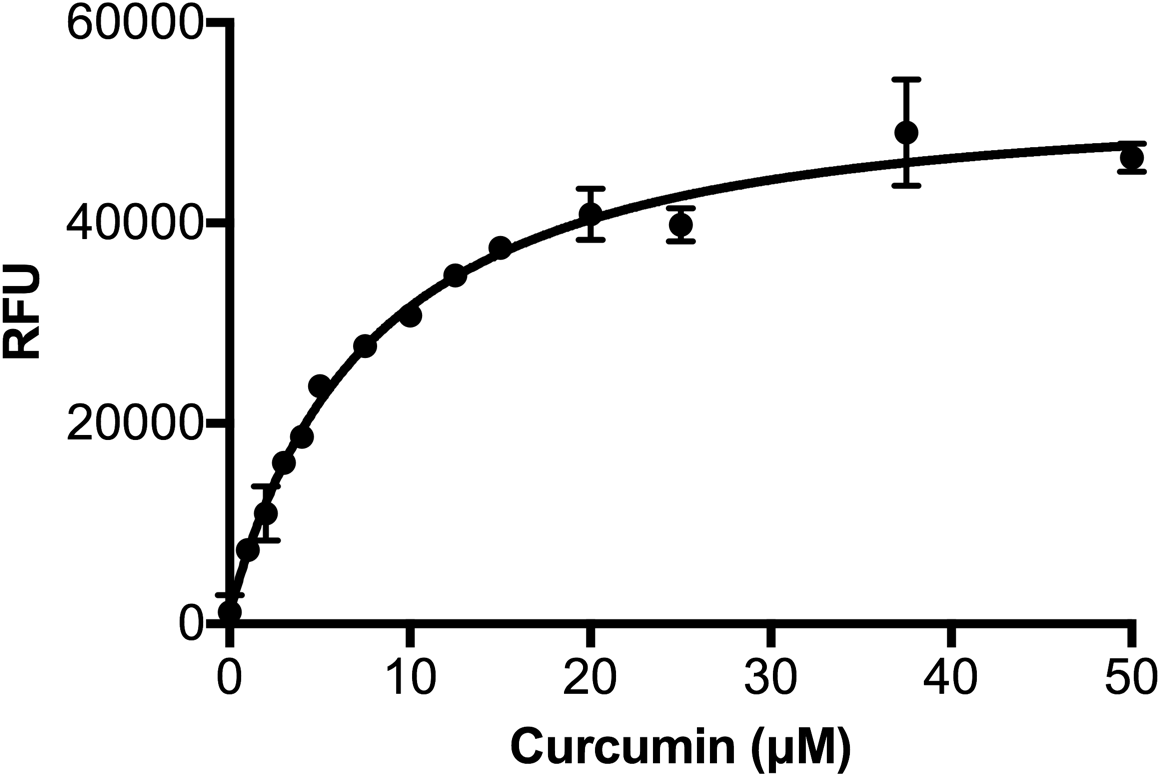
Fluorescence measurement of 25 μM *Ec*CurA with an increasing concentration of curcumin. Data is an average of two biological repeats of purified *Ec*CurA, prepared from separate batches.

**Figure S2.**
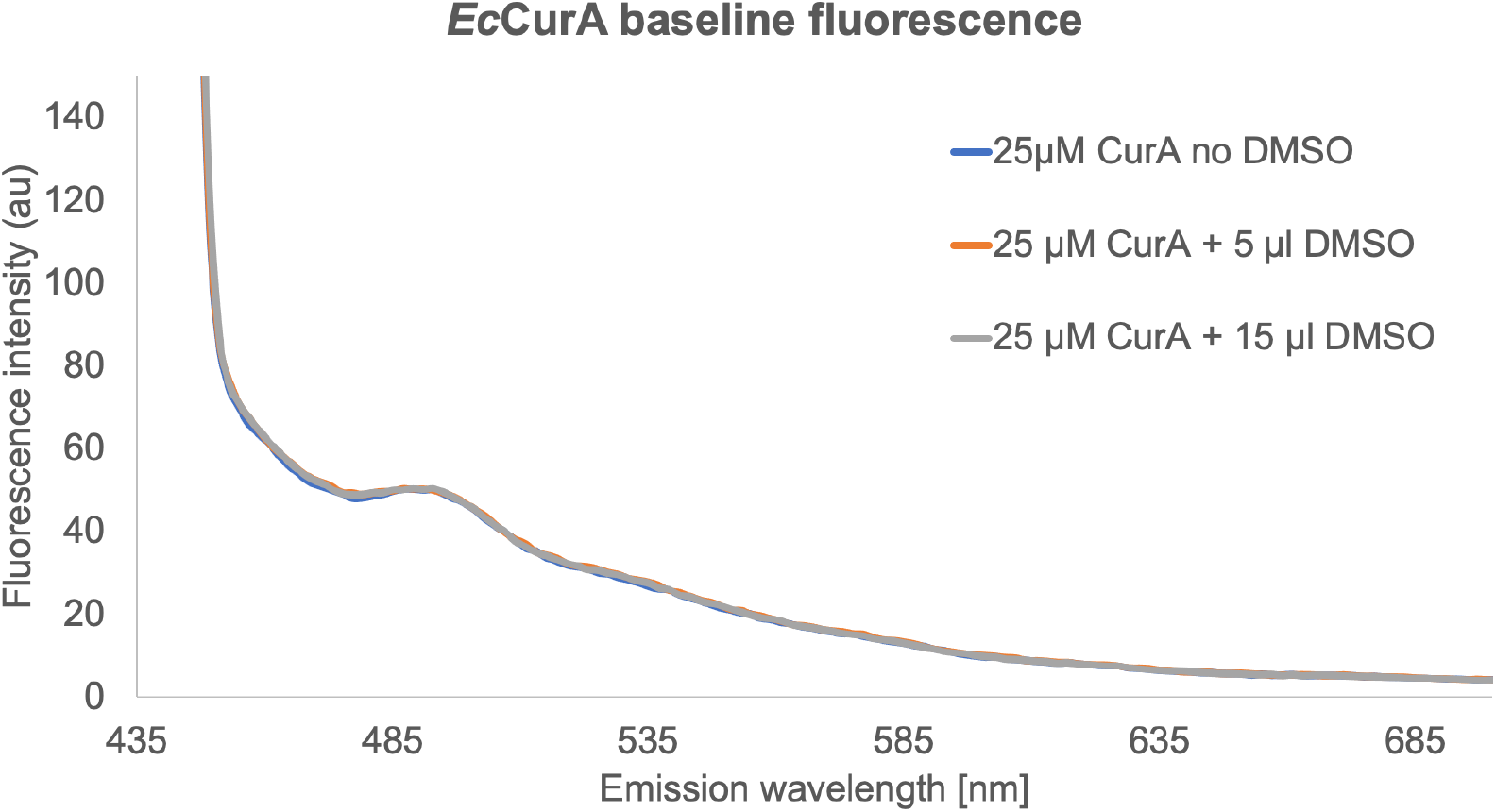
Baseline fluorescence of purified *Ec*CurA. Measurements were prepared in a 3 mL quartz cuvette.

**Figure S3.**
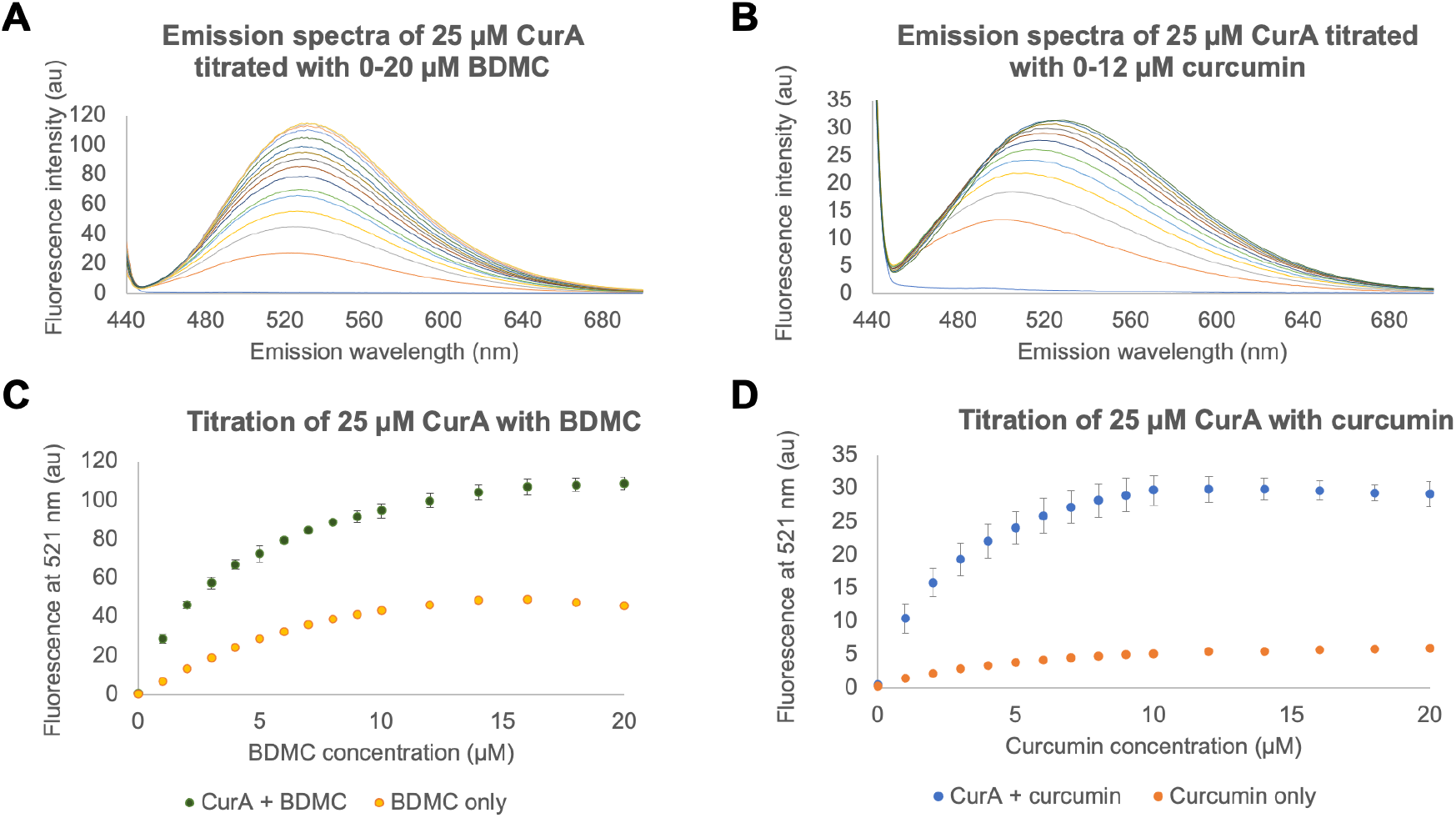
Fluorescence titrations of EcCurA. (A) Emission spectra of *Ec*CurA-BDMC. (B) Emission spectra of *Ec*CurA-curcumin. (**C**) BDMC intrinsic fluorescence spectra in presence or absence of EcCurA. (**D**) Curcumin intrinsic fluorescence spectra in presence or absence of EcCurA. Measurements were prepared in a 3 mL quartz cuvette. Data is an average of two biological repeats of purified *Ec*CurA, prepared from separate batches.

**Figure S4.**
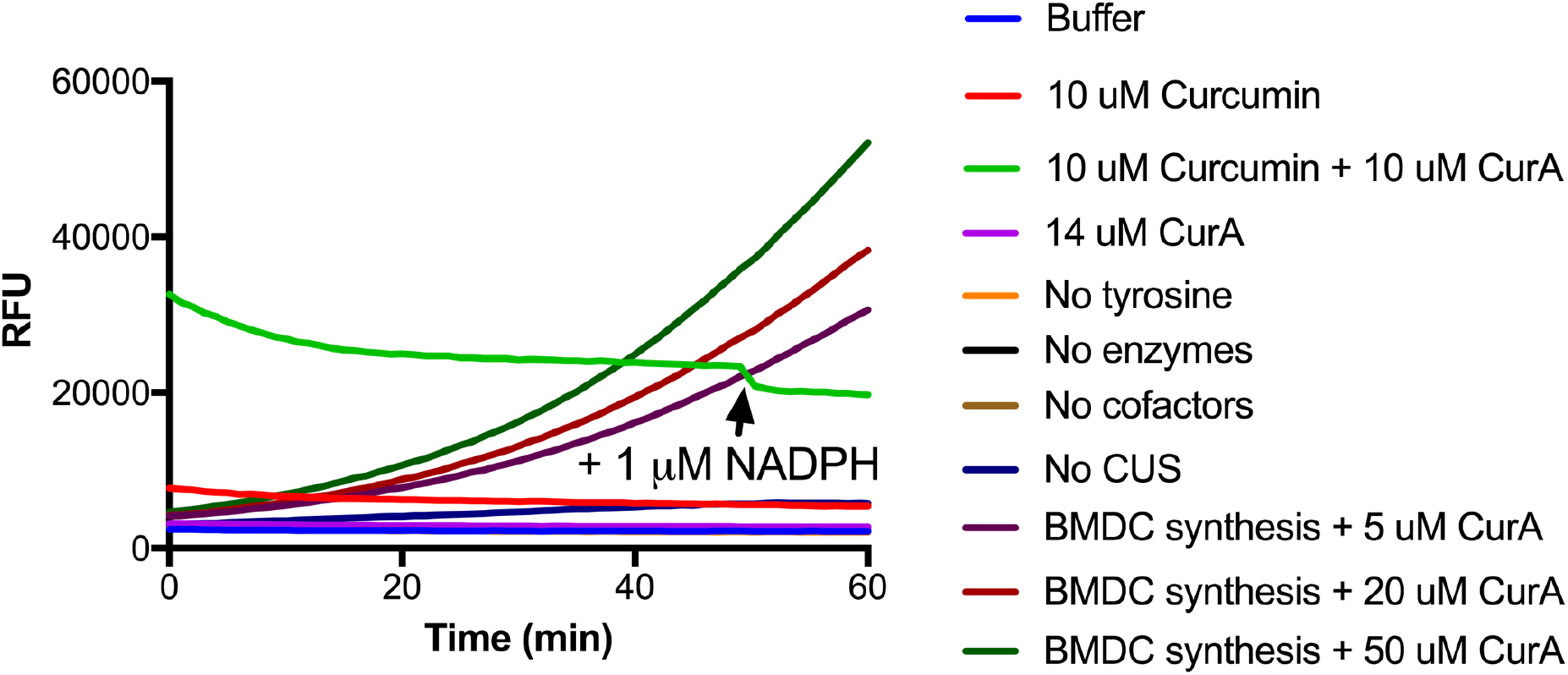
CurA time-course activity assay. An initial assay containing 1 mM L-tyrosine, 5 mM MgCl_2_, 10 mM malonate, 5 mM ATP, 0.25 mM CoA, 5 μM TAL, 5 μM PCL, 5 μM MatB, 5 μM CUS and 25 μM *Ec*CurA. Negative controls are shown whereby individual components of the reaction were omitted, thus showing relative background fluorescence. Unbound BDMC/curcumin has weak intrinsic fluorescence. Injection during the enzyme time-course reaction with 1 μM NADPH is shown by an arrow on the green trace (curcumin and *Ec*CurA). Data is an average of three technical repeats. These reactions are an average of triplicate technical repeat, with error bars removed for clarity.

